# A minimal model of Norway spruce - bark beetle outbreak dynamics

**DOI:** 10.1101/2025.10.27.684373

**Authors:** Alice Doimo, Davide Zanchetta, Giai Petit, Davide Nardi, Andrea Battisti, Sandro Azaele, Amos Maritan

## Abstract

Bark beetles pose a growing threat to coniferous forests worldwide, with outbreaks increasingly linked to environmental stress and host vulnerability. Most mechanistic models focus on equilibrium states, neglecting the dynamical nature of outbreaks triggered by environmental perturbations. Using *Ips typographus* as focal species, we present a minimal model capturing key processes such as threshold-driven mass attacks, with parameters adaptable to different environmental conditions. Under current conditions, the model predicts a stable endemic state that is reactive, meaning environmental shocks trigger isolated, transient outbreaks. Under altered conditions, reflecting potential climate change scenarios, the system undergoes a regime shift, producing recurrent outbreaks that arise spontaneously from insect–host feedbacks. Finally, we examine repeated environmental disturbances and forest management, showing that the effectiveness of outbreak mitigation depends on the interplay between management frequency and intensity. Overall, we provide a mechanistic framework for understanding outbreak dynamics and guiding management strategies under changing environmental conditions.

## I. INTRODUCTION

Conifer-dominated forests constitute a significant component of the boreal and temperate ecosystems throughout Europe and North America [1, 2]. These forests provide essential services such as carbon sequestration, biodiversity preservation, and renewable raw material for economic activities. Within forest environments, biotic agents such as insects and fungi play key ecological roles, influencing nutrient cycling, succession, and structural complexity. Among these biotic agents, bark beetles (Coleoptera Curculionidae Scolytinae) are particularly important due to their capacity to profoundly shape forest structure and successional pathways [3, 4]. Under endemic conditions, bark beetles typically attack and kill weakened or overmature trees, thus contributing to forest regeneration and improving spatial heterogeneity [5, 6]. When population outbreaks occur, tree mortality increases to the point that affected stands may be severely depleted [7–10]. Under certain triggering conditions, bark beetle populations erupt into epidemics that cause severe ecological upheaval. Widespread tree mortality leads to severe forest loss and substantial economic repercussions, impacting timber production and necessitating costly management interventions [11]. Over the past several decades, a variety of modeling approaches has been developed to understand the drivers and dynamics of bark beetle infestations [12, 13]. These range from discrete-time models, often formulated at varying levels of abstraction [14, 15], to continuous-time dynamical systems that aim to capture beetle-host interactions in a more mechanistic framework [16, 17]. While these models yield valuable insights, particularly by identifying stable or unstable equilibria that may govern long-term behavior, they often offer limited understanding of how outbreaks are actually initiated, evolve over time, or recur. In many documented cases, large-scale epidemics do not arise solely from gradual, internal processes operating near equilibrium. Instead, they are often triggered by abrupt external shocks that rapidly alter system conditions. Prominent examples are windstorms, which caused extensive treefall and dramatically increased the availability of vulnerable hosts, leading to a surge in beetle attacks [18–20], and heat waves or drought events that can strongly affect host tree defences [21–23]. Such episodes underscore the fact that outbreaks frequently emerge as transient responses to perturbations, rather than arising naturally from long-term ecological balance, even though climate modification might lead to persistence disturbance regime regionally [24]. Despite ongoing modeling efforts, a mechanistic explanation of outbreak temporal patterns that is both simple and ecologically plausible remains elusive. This gap limits our ability to anticipate or mitigate future outbreaks — especially as climate change increases the likelihood of disruptive environmental events [25–27].

In our work, we address this gap by introducing a minimal dynamical model of bark beetle–tree interactions, using the *Ips typographus* - *Picea abies* system as a case study. Our goal is to explain the outbreak phenomenology of the European spruce bark beetle (ESBB) through a simple mechanistic framework that captures key ecological processes driving outbreak dynamics. Our analysis builds on two complementary perspectives. First, we investigate the stability of typical forest conditions and their reactivity — that is, whether small perturbations are initially amplified or suppressed. Second, we explore the emergence of regularly-recurrent outbreaks arising from ecological feedbacks.

## II. METHODS

### A. Model

We develop a compartmental deterministic model to describe the temporal dynamics of ESBB populations and their interactions with Norway spruce trees within a sufficiently large forest patch, such that spatial heterogeneity and finite-size effects are neglected. The model tracks three state variables over time: the density of bark beetles, denoted by *x*(*t*), and the densities of trees in two distinct physiological states, *healthy y*(*t*) and *vulnerable z*(*t*), the latter signifying immediate susceptibility to beetle attack. Vulnerable trees are intended as those with impaired defenses, typically due to aging, drought/pathogens stress, or mechanical damage (e.g. wind-throw). These individuals are highly susceptible to colonization even at low beetle densities and serve as the primary resource for reproduction. Such distinction is critical for capturing outbreak dynamics, as the presence and abundance of vulnerable hosts strongly influence the potential for beetle population growth and the onset of epidemic phases. The bark beetle population follows consumer–resource dynamics, with individuals dying at rate δand reproducing at a rate *βz*, reflecting the dependence of successful reproduction on access to vulnerable host trees. In addition, beetles arrive via a constant immigration term *λ*, representing an influx of bark beetles from surrounding areas. In real ecosystems, bark beetles disperse through active flight, wind-assisted transport, or anthropogenic movement (e.g., infested wood), giving rise to an immigration pressure from neighboring populations. From a modeling perspective, this term reflects the assumption that the focal system is part of a larger, spatially connected landscape, and prevents unrealistic extinction due to transient local declines, ensuring beetle persistence even under temporarily unfavorable conditions. In the absence of bark beetles, the tree population follows logistic growth with rate *η* and a normalized carrying capacity set to 1. Healthy trees enter the vulnerable state at a baseline rate *ϵ*, representing the effects of aging or moderate environmental stress. Episodic shocks, such as major windstorms or heat waves, are not included as they will be treated as external perturbations (Section III A, Section III D). Vulnerable trees die and are removed from the system at rate *ρ*. The balance between *ϵ* and *ρ* serves as a proxy for forest condition: when trees shift more quickly into vulnerability (high *ϵ*) and are removed more slowly (low *ρ*), the *ϵ/ρ* ratio increases, indicating a more degraded or stressed ecosystem. We model bark beetle–host interactions by incorporating a density threshold mechanism that governs attacks on healthy trees. In the absence of an outbreak, healthy trees become vulnerable at the baseline rate *ϵ*, as described earlier. However, when the beetle density exceeds a critical threshold *σ*, coordinated mass attacks lead to an accelerated conversion of healthy trees into vulnerable ones at rate *κx*. Vulnerable trees, in turn, are depleted as part of the beetle reproductive cycle, with a removal rate *γxz* that reflects their role as the primary substrate for brood development. While beetle aggregation behavior is not explicitly modeled, its effects are implicitly incorporated into the definition of the threshold *σ*, consistent with our mean-field approach [28] as discussed in SI IIA. The system can thus be described at the deterministic level by the following set of coupled differential equations:

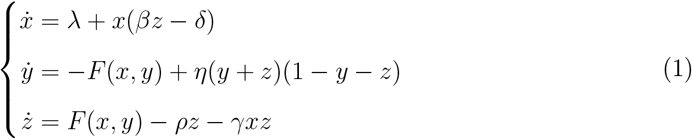

where the function *F* (*x, y*) = *ϵy*+*κ* Θ(*x*−*σ*)*xy* represents the conversion rate from healthy to vulnerable trees, comprising both *background* and *outbreak-induced* conversion from healthy to vulnerable trees, the latter being represented through the Heaviside step function Θ(*x*−*σ*). A derivation from minimal ecological key processes is provided in SI I, together with a schematic representation of the model compartments and transitions.

While the model is simple, capturing only the essential features of bark beetle–host interactions, its core ingredients are sufficient to reproduce key nonlinearities and feedback loops that characterize bark beetle–host dynamics. This balance enables rich ecological phenomenology while retaining analytical tractability. Moreover, the model is flexible, as parameters can be adapted to reflect different forest health conditions or climate scenarios.

### B. Parameter assumptions and estimation

We adopt a set of simplifying assumptions that specify how beetle and tree processes are represented. These abstractions are coarse but crucial for tractability. In particular, although bark beetles reproduce in discrete generations and are largely inactive during winter, we model processes by averaging seasonal dynamics over the year. Since reproduction depends on ovipositing females, we track only the adult female population. Accordingly, we model recruitment as half of the total brood size, assuming a 1:1 sex ratio, while juvenile stages are not explicitly represented. Trees smaller than the minimum size required for successful attack by ESBB are not explicitly tracked. Tree population growth is modeled with a logistic function, reproducing both host replenishment and stand saturation. Mass attacks on healthy trees require a critical number 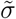 of insects per tree. ESBB are known to aggregate via pheromone chemical signaling over a finite spatial range. As a result, each tree is exposed to ESBB within its neighborhood, and we accordingly assume that mass attacks are triggered when the number of insects within this aggregation area exceeds 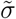. While the threshold *σ* captures the resistance limit of trees, the colonization rate *κ* describes how efficiently insects exploit that vulnerability under outbreak conditions. Since no direct empirical estimates are available, *κ* is treated as a free parameter in our scenario analysis.

Based on these assumptions, we estimate order-of-magnitude values for all parameters from the ecological and entomological literature. These values are not tuned to any specific local ecosystem but are intended to reflect typical conditions of Norway spruce forests in Europe. This level of approximation is sufficient to investigate the model’s qualitative dynamics, such as the emergence of equilibria or recurrent outbreaks. However, more precise estimates—tailored to specific ecosystems—would be required for reliable quantitative predictions (e.g. outbreak duration or magnitude). Table I summarizes the adopted reference values and plausible variability ranges. The latter reflect both ecological heterogeneity (e.g., altitude, forest density) and measurement uncertainty, and they define what we refer to as the ecologically plausible region of parameter space. Further details on parameter estimation are provided in SI II. While the values presented here provide a consistent reference framework for analyzing the model’s behavior, they can readily be adjusted to reflect different ecological or forest structural conditions.

**TABLE 1.** Reference values and variability ranges for the model’s parameters, representative of generic present-day environmental conditions. We estimate reference values for the model’s parameters based on available ecological and entomological literature [4, 29–34]. These estimates reflect current typical conditions in managed European spruce forests, while also accounting for known sources of variability, derived from ecological heterogeneity or uncertainty in empirical measurements. Where precise data are lacking, we use informed assumptions to define plausible orders of magnitude or leave parameters free for exploration. All details on parameter meaning and estimation are provided in SI II, together with the references to the literature.

| Parameter | Description | Reference value | Variability range |
| --- | --- | --- | --- |
| $L^2$ | Area of the forest patch | 1 ha | fixed |
| $M$ | Maximum number of trees per patch | 500 | 300 – 700 |
| $\rho$ | Death rate of vulnerable trees | $0.001 \text{ day}^{-1}$ | $[6 \cdot 10^{-4}, 5 \cdot 10^{-3}] \text{ day}^{-1}$ |
| $\epsilon$ | Conversion rate (healthy → vulnerable) | $1 \cdot 10^{-5} \text{ day}^{-1}$ | $[7 \cdot 10^{-6}, 3 \cdot 10^{-5}] \text{ day}^{-1}$ |
| $\eta$ | Birth rate of healthy trees | $1.3 \cdot 10^{-2} \text{ day}^{-1}$ | $[4 \cdot 10^{-3}, 3 \cdot 10^{-2}] \text{ day}^{-1}$ |
| $\delta$ | Death rate of ESBB | $0.5 \text{ day}^{-1}$ | $[0.3, 1.0] \text{ day}^{-1}$ |
| $n$ | Number of insects per brood (females) | 25 | 20 – 30 |
| $\beta$ | Reproduction rate of ESBB | $18 \text{ day}^{-1}$ | $[5, 40] \text{ day}^{-1}$ |
| $\gamma$ | Tree death rate due to beetles | $0.4 \text{ day}^{-1}$ | $[0.1, 0.7] \text{ day}^{-1}$ |
| $\sigma$ | ESBB density threshold for mass attack | 10 | 3 – 85 |
| $\kappa$ | Colonization rate of healthy trees | Free | – |

### C. Equilibria

The first step in analyzing dynamical models is to identify their equilibria — population levels at which ESBB and trees remain approximately constant over time. Ecologically, these represent long-term configurations of population stability, without rapid growth or collapse, maintained by a balance among interacting biological processes. These states are stable, meaning that if an environmental disturbance briefly perturbs the system, populations will eventually return to their previous levels.

The following theoretical classification outlines the types of steady states that the system can, in principle, sustain under different climatic scenarios, as determined through mathematical analysis (SI III). Depending on parameter values, the model admits qualitatively distinct equilibria:

i. The *trivial equilibrium* describes a forest in which healthy and vulnerable trees are absent, and the ESBB population persists only through external immigration. Without suitable hosts, beetles cannot reproduce, and in the absence of immigration (*λ* = 0) they decline to extinction. This state is ecologically meaningless, as neither trees nor a self-sustaining beetle population can persist.
ii. The *endemic equilibrium* is a state where both trees and ESBB persist, but beetle density remains too low to trigger mass attacks on healthy trees, representing a forest that is not inherently prone to outbreaks. The existence of this stable state requires immigration (*λ >* 0): without it, beetles would go extinct, yet even minimal influx is sufficient to maintain a low-density population.
iii. The *epidemic equilibrium* is characterised by an ESBB abundance high enough to mount mass attacks on healthy trees. In this state, the ESBB remains permanently above the mass-attack threshold, corresponding to an indefinitely sustained outbreak.
iv. Lastly, *out-of equilibrium regimes* occur under climatic conditions where no stable equilibrium exists. In this case, ESBB and tree populations fluctuate persistently, with outbreak and forest recovery phases that never settle into a steady state.

Although identifying stable equilibria provides insight into the long-term baseline behavior of the ecosystem, it is not sufficient to capture outbreak dynamics. Within this equilibrium framework, the system can only display three idealized behaviors: a complete absence of outbreaks (endemic state), a permanently sustained outbreak (epidemic state), or endless collapse and recovery (out-of-equilibrium regime). None of these configurations reflects the typical forest dynamics observed in reality, where outbreaks are transient events triggered by environmental disturbances. To address this limitation, we extend the analysis to dynamical properties beyond equilibrium stability and we examine *reactivity*, which quantifies the system’s short-term response to perturbations.

### D. Reactivity

To characterize outbreak dynamics beyond long-term equilibria, we analyze the system’s reactivity [35], which quantifies the short-term response of a system in a stable equilibrium state, when it is perturbed e.g. by environmental disturbances. Unlike classical equilibrium stability analysis, which assesses only whether small perturbations eventually decay, reactivity captures whether they temporarily grow before decaying. This transient amplification leads to short-lived outbreaks, even though the system ultimately returns to equilibrium.

Mathematically, reactivity is assessed via the symmetric part of the Jacobian matrix 𝕁 evaluated at the equilibrium:

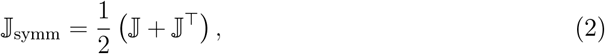

Letting *λ*_1_, *λ*_2_, *λ*_3_ be the eigenvalues of 𝕁 _symm_, we define reactivity as:

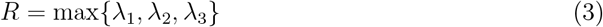

The system is reactive if *R >* 0, meaning disturbances temporarily grow before decaying, causing an outbreak; otherwise (*R <* 0) it is non-reactive and perturbations decay monotonically. Importantly, when positive, *R* provides an intrinsic measure of outbreak propensity: the larger its value, the stronger the amplification of a given environmental disturbance, leading to more intense transient outbreaks.

## III. RESULTS

Our analysis builds on two complementary dynamical aspects introduced earlier: (i) reactivity, which characterizes the system’s transient response to small perturbations near equilibrium, and (ii) intrinsic regularly recurring outbreaks emerging from nonlinear ecological interactions.

### A. Transient outbreaks triggered by disturbances

We assess the reactivity of the endemic equilibrium state, representing the most ecologically plausible scenario under typical conditions (Table I) for managed European spruce forests. This equilibrium is consistent with observations from undisturbed stands where beetles persist at low-densities, insufficient to trigger mass attacks. Despite the structural stability of the endemic equilibrium, outbreaks commonly occur in real forests. This apparent contradiction is explained by the reactivity of the endemic equilibrium under realistic conditions. As explained in Section II D, a positive eigenvalue of the symmetrized Jacobian J_symm_ indicates reactivity, i.e. small perturbations of the equilibrium state of the forest ecosystem initially grow, before eventually decaying. In contrast, if all eigenvalues are negative, the equilibrium is non-reactive, and all perturbations decay monotonically.

As shown in Fig. 1(a), the most reactive perturbation (along the eigenvector of the positive eigenvalue) leads to a transient spike away from equilibrium, whereas the two nonreactive ones (related to negative eigenvalues) yield monotonic relaxation. The structure of the eigenvectors (SI IV) reveals that this transient growth is primarily driven by changes in the susceptible tree population. In contrast, perturbations along non-reactive eigenvectors are dominated by changes in healthy trees or beetles alone, and do not produce significant transient growth. This suggests that reactivity in the ESBB model is mainly driven by perturbations that abruptly increase the abundance of vulnerable trees, such as extreme windstorms or heat waves. To illustrate reactive behavior, Fig. 1b) shows population trajectories following a perturbation strongly aligned with the reactive eigenvector. The ESBB population exhibits a sharp but temporary increase before returning to equilibrium, capturing the hallmark of a transient outbreak. The duration of the outbreak and the height of the ESBB peak depend on two factors: first, the reactivity index *R*, which reflects the background environmental conditions within the typical parameter range (Table I); second, the strength of the perturbation, representing the environmental shock.

**FIG. 1.**
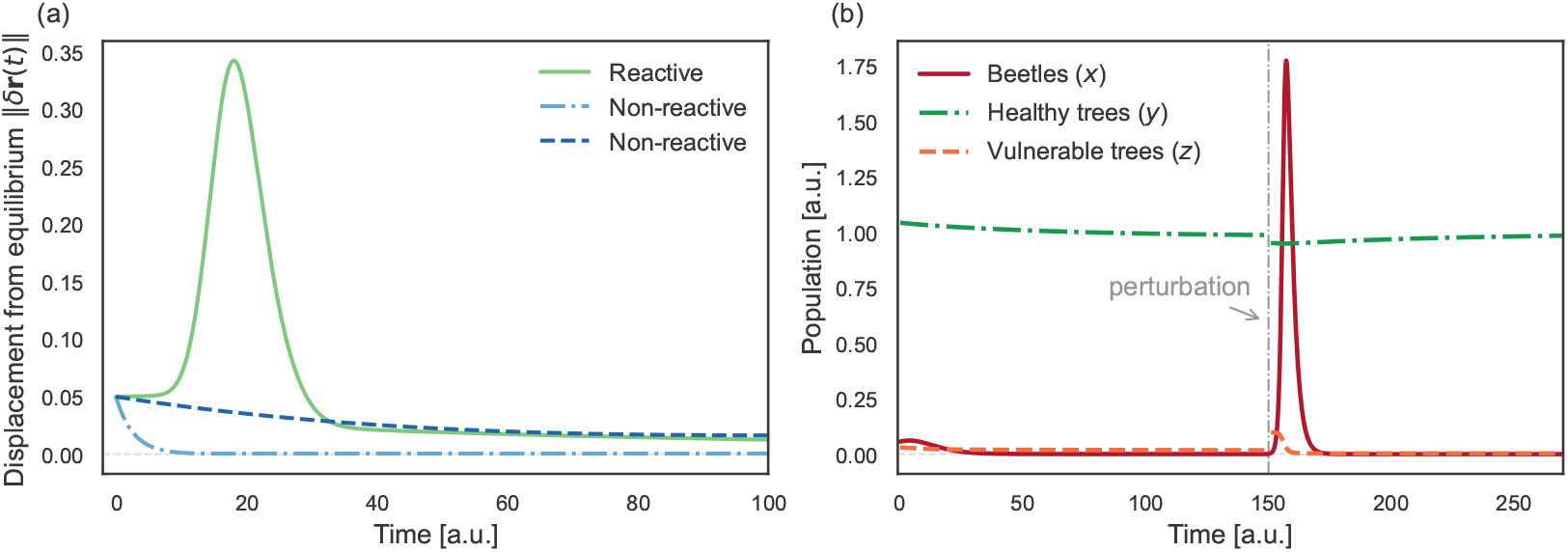
Reactive dynamics in the ESBB model. **(a)** System response to small perturbations applied at *t* = 0 along different directions defined by the eigenvectors of the symmetrized Jacobian at equilibrium. A perturbation along the most reactive direction (green) causes transient amplification of the distance from equilibrium before decay — characteristic of a reactive system. In contrast, a perturbation along non-reactive directions (blue) decays monotonically. **(b)** Time evolution of the full system. After reaching equilibrium, a small perturbation is applied at *t* = 20 along a direction close to **v**_1_ (*ε* =0.1, δ**r**(*t*)· **v**_1_ ∼ 0.89). This causes a sharp, temporary increase in beetle population (red), before the system returns to equilibrium. Healthy trees (green) decline slightly, while vulnerable trees (orange) show a brief increase. The system’s strong transient response reflects its reactivity. All quantities are given in arbitrary units (a.u.).

We emphasize that, although the ESBB population may temporarily exceed the massattack threshold during the transient growth phase, this increase is short-lived because the conditions required to sustain epidemic activity are not met, and the system eventually relaxes back to the sub-threshold endemic state. If epidemic conditions were fulfilled, by contrast, the system would settle into an epidemic equilibrium, where ESBB abundance would remain permanently above the mass-attack threshold, corresponding to a persistent outbreak state. Fig. 2(left) shows that reactivity exhibits a dramatic, discontinuous jump as the system transitions from the endemic to the epidemic equilibrium, and reveals a link between equilibrium beetle density and outbreak propensity.

**FIG. 2.**
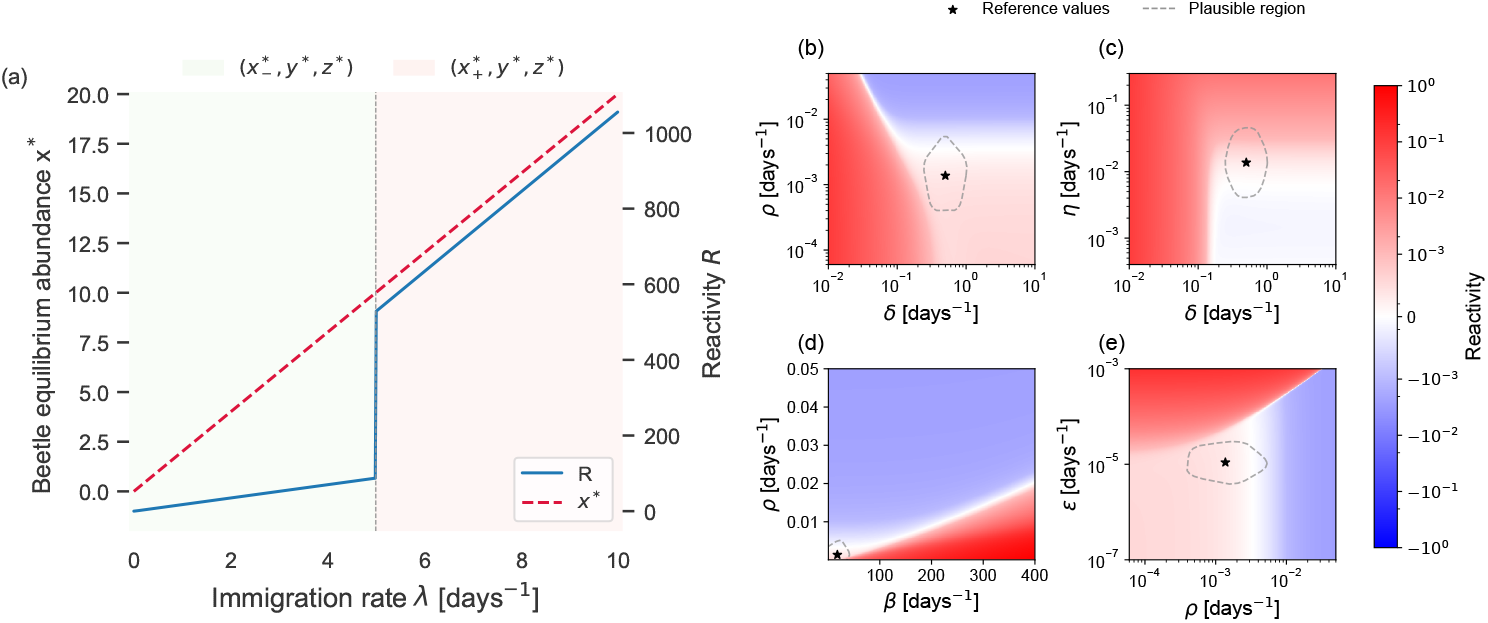
(left) Reactivity and beetle equilibrium abundance as a function of the immigration rate parameter *λ*. As *λ* increases, the beetle equilibrium abundance grows linearly. When *x*^∗^ crosses the mass attack threshold *σ* = 10, the system transitions from a regime characterized by the stable endemic equilibrium (*x*^∗^_−_, *y*^∗^, *z*^∗^) to a regime where the above-threshold fixed point (*x*^∗^_+_, *y*^∗^, *z*^∗^) becomes stable. Both regimes are reactive and display a linear increase of reactivity with *λ*, indicating growing sensitivity to perturbations at higher immigration rates. Notably, at the critical immigration rate (*λ* = 5), reactivity undergoes a discontinuous jump in value, accompanied by a sharp change in slope, reflecting a fundamental shift in system dynamics upon crossing the outbreak threshold. The remaining parameters are kept fixed at the values reported in Table I. **(right) Reactivity landscape**. Reactivity over a region of parameter space centered on the reference values given in Table I. The star marks the reference parameter set, at which the reactivity is *R* ≈ 0.002. As one moves into the ecologically plausible regime surrounding the reference point, which represents approximate but realistic ranges for model parameters, the system becomes consistently reactive. Thus, despite uncertainties in precise parameter values, the system is generally expected to exhibit reactive dynamics under realistic ecological conditions, suggesting inherent susceptibility to transient outbreaks following perturbations.

### B. Phenomenology of dynamical behaviors across ecological conditions

We use *phase diagrams* to explore how ecological parameters shape ecosystem dynamics. These diagrams show which equilibria arise under different conditions, capturing long-term system behavior. Complementarily, we map the reactivity of each equilibrium to assess short-term sensitivity to perturbations. Here, two parameters are varied along the axes, while color coding indicates equilibrium type, providing a compact view of how ecological conditions can shift the system from endemic stability to more outbreak-prone or unstable regimes.

Fig. 3(a) illustrates the model’s dynamical richness when parameters are broadly varied beyond ecologically grounded values reflecting current forest conditions (Table I). Fig. 3(c) shows representative time trajectories illustrating the range of possible behaviors under different parameter combinations. These trajectories are not intended to reproduce realistic time series for specific ecosystems, nor to predict the exact timing, duration, or magnitude of real outbreaks. Rather, they illustrate possible outcomes under different ecological scenarios, represented here by varying the underlying parameters. We observe: (i) monostable regimes (green, blue, and red) where populations approach unique equilibria, potentially after transient oscillations; (ii) bistable regimes (yellow and violet) where the attained state sensitively depends on initial conditions, resulting in alternative stable attractors; and (iii) regions lacking stable equilibria (grey), where populations exhibit regularly recurrent, selfsustained outbreaks. The latter regime is examined in depth in Section III C. Importantly, in highly reactive endemic equilibria with complex eigenvalues (as in the blue panel), oscillations that cross the mass-attack threshold *σ* can trap the system in recurrent outbreak dynamics, even though the equilibrium is formally stable.

**FIG. 3.**
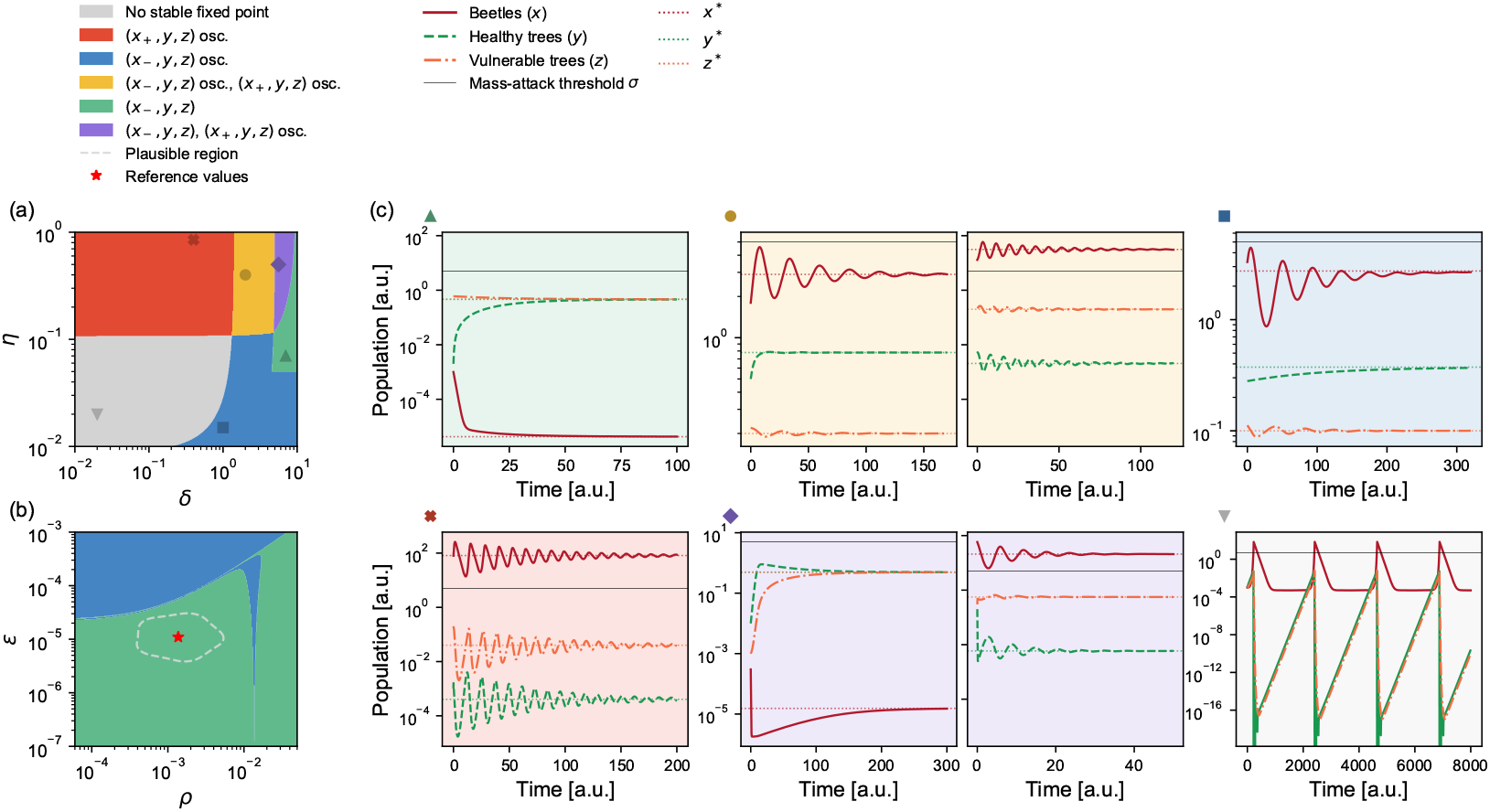
Phenomenology of the model. **(a)** Phase diagram computed over a wide, arbitrary parameter range to illustrate the diversity of possible dynamical behaviors supported by the model, including stable coexistence, oscillations, and instability. These values are not ecologically grounded but demonstrate the system’s inherent dynamical richness. Parameters *ϵ* and *κ* are varied, while all others are held fixed at the values reported in SI VIII. **(b)** Phase diagram restricted to broad parameter ranges centered around the ecologically grounded reference values reported in Table I. The highlighted region approximately denotes an plausible ecological regime, representative of managed European spruce forests. Even well beyond this region, the system consistently converges to a stable endemic equilibrium characterized by sub-threshold beetle densities. **(c)** Trajectories of beetle, healthy, and vulnerable tree populations for parameter values sampled from distinct regions of the phase diagram in Fig. 3(a), where the corresponding points are marked with diamond symbols. Each panel corresponds to a qualitatively different dynamical regime, that is not intended as a realistic scenario, but rather as a qualitative exploration of the different possible behaviors of the system over time. In bistable regions (yellow and violet), the system can converge to different stable fixed points depending on the initial conditions. Notably, in regions where no stable fixed point exists (grey phase), we consistently observe sustained oscillations, indicative of limit cycles, which will be analyzed in detail in Section III C. The labels *x*^∗^, *y*^∗^, *z*^∗^ indicate the equilibrium values for ESBB, healthy and vulnerable tree populations, respectively. We remark that population abundances and timescales are given in arbitrary units (a.u.).

Fig. 3(b) focuses on ecologically plausible parameter ranges for managed European spruce forests (Table I). Within this region, the system robustly settles into a reactive endemic equilibrium (with non-complex eigenvalues), discussed in Section III A. Notably, the endemic equilibrium remains robust across a broad parameter region, suggesting resilience beyond finely tuned ecological conditions.

Fig. 2 visualizes reactivity’s dependence on ecological parameters. Outside realistic parameter ranges (Table I), the endemic equilibrium is typically non-reactive (blue), meaning disturbances decay directly without causing outbreaks. Within ecologically grounded values, the system becomes consistently reactive (red), indicating high sensitivity to small shocks that result in outbreaks. As expected, reactivity increases with longer adult beetle flight period δ (panels a-b) or higher ESBB reproductive rate *β* (panel c), while faster removal of vulnerable trees (higher *ρ* in panels c-d) reduces it. An elevated tree susceptibility *ϵ* also raises reactivity (panel d), linking increased environmental stress to higher transient outbreak risk. Additional reactivity maps for other parameter pairs are shown in Fig. S2.

### C. Recurrent outbreak dynamics

As noted in Section II C, for some parameter choices outside the reference range representing current environmental conditions, the model does not admit any stable equilibrium. In such cases, there are in principle three possibilities: populations may diverge, fluctuate chaotically, or oscillate regularly. In our model, we observe only the latter. Such oscillations are known as *limit cycles* in dynamical systems theory, but for simplicity we refer to them here as recurrent outbreaks. Interestingly, such regularly-recurring outbreaks also emerge in parameter regions where a stable equilibrium exists but is effectively inaccessible due to extreme reactivity: even minimal increases in the number of susceptible trees can push the system away from equilibrium and trigger recurrent outbreaks (Fig. 4(a).

**FIG. 4.**
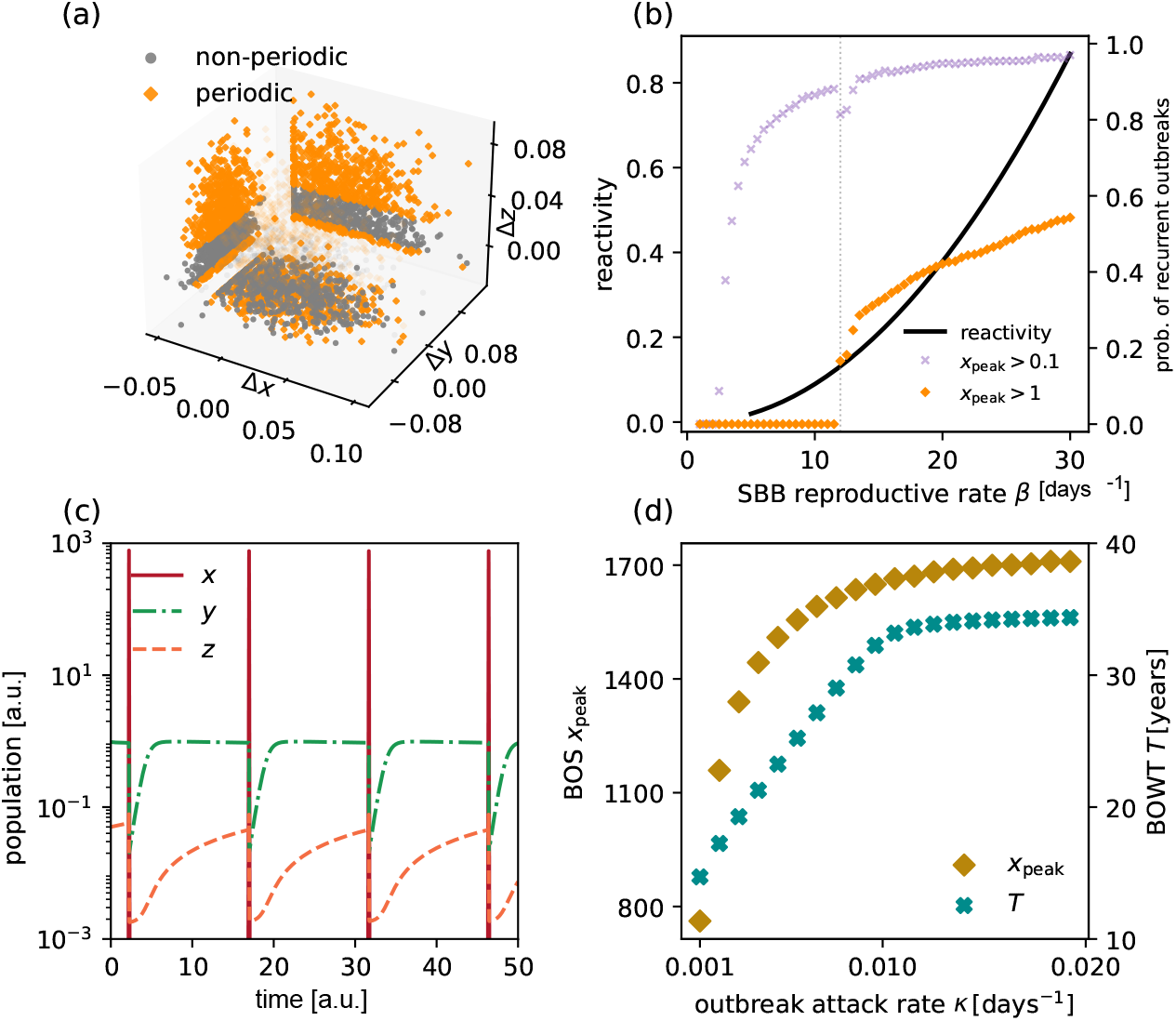
Emergence of recurrent outbreaks with increasing reactivity. **(a)** For a fixed set of parameters, we perturb equilibrium with a Gaussian vector. We project the ensemble of 10^4^ sampled perturbations, (Δ*x*, Δ*y*, Δ*z*), on the coordinate planes. **(b)** Varying *β* causes reactivity of the endemic equilibrium to grow continuously (solid line). Instead, the probability of observing recurrent outbreaks (*x*_peak_ *> x*_thr_ with *x*_thr_ ⪆ 1) increases sharply at *β* ≈ 12 (dotted vertical line), indicating a new dynamical phase. Detecting recurrent dynamics with a smaller threshold (*x*_peak_ *> x*_thr_ = 0.1) gives qualitatively consistent results over a broad range of values for the threshold, while also counting small decaying oscillations. gives qualitatively consistent results over a broad range of values for the threshold, while also counting small decaying oscillations. As in panel (a), outbreak probability has been computed by sampling initial conditions for Eq. (1) ffrom a Gaussian neighborhood of the stable equilibrium value and automatically detecting recurrent outbreak dynamics. **(c)** Population dynamics for ESBB *x*, healthy trees *y* and susceptible trees *z* (densities). The peaks of ESBB population *x*(*t*) define the baseline outbreak size (BOS), while their temporal spacing defines the baseline outbreak waiting time (BOWT). Population abundances and timescales are given in arbitrary units (a.u.). **(d)** Outbreaks observables: baseline outbreak size *x*_peak_ and baseline outbreak waiting time *T* for different values of *κ*. Note that the period and intensity of outbreaks are sensitive to parameter choices; since ecological parameters are not precisely constrained, these results should be interpreted as a phenomenological investigation of the model’s dynamical behavior rather than quantitative predictions of BOS and BOWT. Full parameter values and numerical details are reported in SI VIII.

Recurrent outbreak dynamics arise intrinsically from internal ecological feedbacks and interactions. In the time interval between outbreaks, the ESBB population remains close to zero, and healthy and susceptible trees tend towards a steady balance. As susceptible trees accumulate, the insect population starts growing exponentially, even from very low densities, provided it is not exactly zero. Once they exceed the density threshold *σ*, insects begin attacking healthy trees, which further accelerates their growth. As host trees are quickly depleted, insect population soon recedes to endemic levels, allowing the forest stand to recover through tree regrowth and the cycle to begin again (Fig. 4(c).

To study the recurrent outbreak regime, we select parameter values that lie outside the ecologically plausible range representing current forest conditions (Table I), but still preserve a realistic balance between ecological rates. This parameter region is not intended to reflect present-day dynamics, but rather a potential future state of the system under increased ecological stress — for example due to climate change or other long-term disturbances. In particular, we increase the *ϵ/ρ* ratio with respect to the current reference condition (Table I), implying that trees remain vulnerable for longer on average, thereby reflecting increased forest stress. More details are available in SI V.

We characterize recurrent outbreaks using two key observables: the maximum population size during an outbreak (*baseline outbreak size* BOS), and the period between successive outbreaks (*baseline outbreak waiting time* BOWT). These observables allow us to investigate how outbreak intensity and frequency respond to changes in ecological parameters.

As shown in Fig. 4b, boom-and-bust dynamics become more likely as reactivity increases — for instance when the ESBB reproduction rate *β* is higher. Another key parameter is the attack efficiency on healthy trees, *κ*, which remains unconstrained by available data. We find that both BOS and BOWT increase with *κ* up to a saturation point (Fig. 4(d). However, recurrent outbreaks are sustained only within an intermediate range of *κ* values: if *κ* is too low, beetles can’t attack healthy hosts; if too high, host trees are depleted during the first outbreak, preventing long-term persistence of the insect population. Despite the uncertainty in its estimation, recurrent outbreaks emerge robustly across a broad range of *κ* values, highlighting the intrinsic capacity of the system to generate recurrent outbreaks through internal ecological feedbacks alone.

### D. Repeated environmental disturbances

We investigate repeated disturbances under the recurrent outbreak regime, which we adopt as a plausible climate-change scenario where forests become intrinsically more vulnerable to pest outbreaks. In addition, we include extrinsic disturbance events — such as heat waves, droughts and windstorms [36] — modeled as sequences of shocks that temporarily increase forest susceptibility to ESBB attacks. While Section III A focused on the endemic state, which reflects current conditions, and examined the effect of a single isolated disturbance, the analysis here reflects the expectation that under climate change endemic coexistence may disappear, recurrent outbreaks prevail, and environmental shocks become more frequent rather than isolated. Each disturbance converts a fraction of healthy trees into susceptible trees, representing the weakening of host defenses. Importantly, we do not model isolated shocks, but a repeated sequence of disturbances over time, mimicking the recurrent stress episodes that forests may experience under changing environmental conditions. Disturbances occur at random intervals of time and with varying intensity, allowing us to explore a broad range of environmental scenarios. Higher frequency represents increasingly unstable climate conditions, while greater severity corresponds to more intense stress events, such as severe droughts. The effect of these disturbances is to modify the otherwise regular dynamics of recurrent outbreaks by introducing variability in both their timing (BOWT) and size (BOS). As shown in Fig. 5a-f, more frequent or severe disturbances lead to smaller but more frequent outbreaks, although rare large outbreaks still occur. Mathematical details of the disturbance implementation are provided in SI VI.

**FIG. 5.**
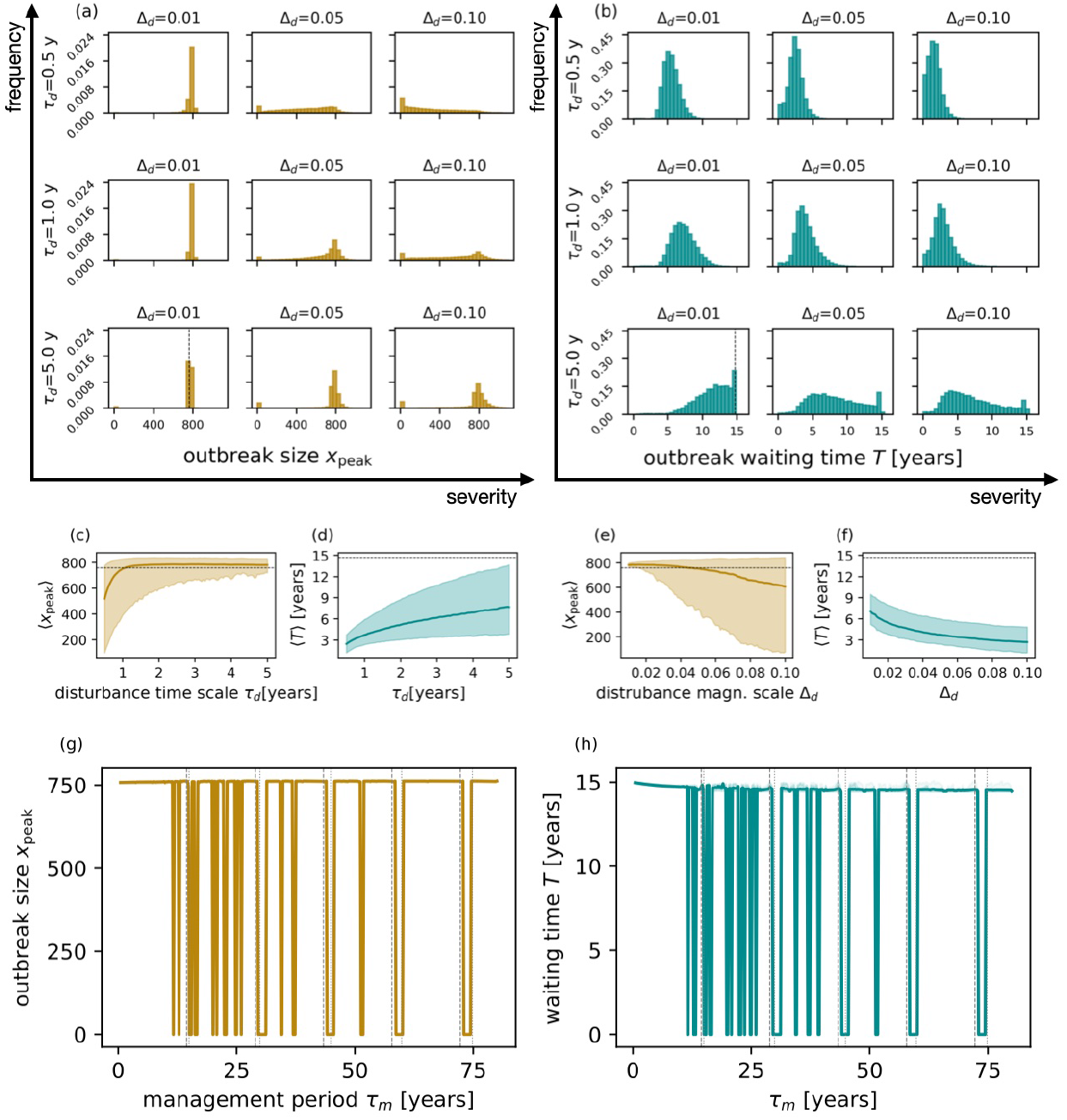
Outbreaks’ size and waiting times change under different disturbance and management regimes. **(Top)** Empirical distribution of outbreak size *x*_peak_ and waiting time *T* for different disturbance parameters *τ*_*d*_ and Δ_*d*_,defining the scales of the exponential distributions of the random size and time of the disturbances. (a-b) Distributions of outbreak size *x*_peak_ (a) and waiting time *T* (b) for different choices of perturbation time scale *τ*_*d*_ and magnitude scale Δ_*d*_. (c-e) Mode (solid line) and 10-90 quantiles (envelope) for *x*_peak_ and *T*, for fixed Δ_*d*_ 0.05 and different values of *τ*_*d*_ (c-d), and for fixed *τ*_*d*_ = 1 year and different values of Δ_*d*_ (e-f). All dashed lines represent BOS and BOWT of the corresponding unperturbed dynamics. **(Bottom)** Outbreak size *x*_peak_ (g) and waiting time *T* (h) as functions of the management periodicity *τ*_*m*_, in a regime of mild disturbances for which ⟨*x*_peak_⟩ ≈ 750 and ⟨*T*⟩ ≈ 14.97 without management; in the presence of management, when outbreaks are not avoided, their mean waiting time decreases significantly, with ⟨*T* ^*′*^⟩ ≈ 14.45 at *τ*_*m*_ = 80. Only *x*_peak_ *> σ* have been counted, with *T* = 0 corresponding to absence of outbreaks. Solid lines are means and envelopes are 10-90 and 20-80 quantiles. Dashed and dotted vertical lines mark values of *τ*_*m*_ which are integer multiples of ⟨*T* ^*′*^⟩ and ⟨*T*⟩, respectively; we note that the former values consistently signal effective management timing. Results are obtained with interdependent Δ_*m*_ and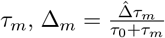. Statistics are accumulated over 1000 realizations for each value of *τ*_*m*_. Timescales and peaks are given in arbitrary units (a.u.). Further details are reported in SI VIII.

### E. Management strategies

We adopt the recurrent outbreak regime as a future scenario to investigate how forest management strategies might mitigate outbreak dynamics. In this context, we represent management as discrete removal events that reduce the density of susceptible trees at regular intervals — an idealized approximation of management practices aimed at lowering stand vulnerability. Although forest operations typically remove both vulnerable and healthy trees, in the model only susceptible trees are removed. This simplification is justified because our analysis shows that system reactivity — and thus outbreak response — depends almost exclusively on the density of susceptible hosts. As a result, selective removal in the model effectively reproduces the ecological outcome of unselective thinning, while allowing a more transparent mechanistic interpretation.

Specifically, we model management as the removal of a fixed fraction Δ_*m*_ of susceptible trees every *τ*_*m*_ time units, as detailed in SI VII. We then explore how the frequency and intensity of removal influence outbreak dynamics. If interventions are too frequent, each action removes only a very small fraction of vulnerable trees, making it ineffective unless the removal rate is unrealistically high. Conversely, if interventions are too rare, the ESBB population may already reach outbreak levels before the next removal event takes place. Crucially, we find that outbreaks can be reliably suppressed when management intervals are tuned to be slightly longer than the system’s natural recurrence timescale (Fig. 5).

## IV. DISCUSSION AND CONCLUSIONS

Bark beetles such as *Ips typographus* play a key role in forest ecosystems, by removing vulnerable trees and accelerating decomposition processes, thereby enhancing the overall heterogeneity [37, 38]. However, under certain conditions, their activity can shift from background ecological function to large-scale disturbances, leading to widespread tree mortality.

Mechanistic descriptions of beetle–forest interactions [16, 17] have so far been restricted to static endemic or epidemic states, thus neglecting the dynamical nature of outbreaks triggered by ecological perturbations. In contrast, process-based, detail-rich models (e.g. [33, 36, 39, 40]) have examined outbreak drivers in depth, but often at the expense of analytical tractability. By reducing the system to its essential components, minimal models clarify the conditions under which forests shift between stability, outbreaks, or collapse. Rather than replicating site-specific details, such models highlight general principles, offering insights that complement more detailed approaches and serving as a foundation for ecological theory and the anticipation of future outcomes. Yet a comprehensive understanding of beetle–forest outbreaks as dynamical events within a mechanistic framework is still lacking. In this study we address these limitations by adopting a dynamical systems perspective, treating outbreaks as transient phenomena driven by ecological feedbacks. Our minimal mechanistic model of beetle–host interactions captures how isolated or regularly-recurring outbreaks can emerge from the interplay of ecological rates such as reproduction, mortality, and host susceptibility. It is important to clarify the interpretative scope of our results. We do not aim to precisely reproduce the exact timing or amplitude of observed outbreaks in a given forest, which would require detailed empirical data and site-specific calibration. Rather, our goal is to identify general dynamical regimes that forest systems may enter under different environmental conditions, represented by varying ecological parameters.

Under current ecological conditions, parameterized using empirically grounded values from spruce–beetle ecology, our model predicts a reactive endemic equilibrium: the bark beetle population remains at low levels, but perturbations can trigger transient outbreaks before the system eventually returns to equilibrium. This behavior is consistent with the available literature, which indicates a low population density during endemic phases and high density and tree mortality during the outbreak phases [4, 41]. Outbreak propensity, captured by the reactivity index, quantifies how strongly a system amplifies small perturbations. Outbreak intensity varies with background environmental conditions and with the strength of the triggering shock. Notably, changes in the abundance of susceptible trees emerge as the primary driver of reactive dynamics. Again, our model captures the effect of the most common triggering factors, such as windstorms and heat waves, which suddenly increase the amount of vulnerable trees [42, 43], facilitating the beetle reproduction and thus sustaining the build-up of the epidemic phase.

We next investigate the model’s predictions when ecological parameters are shifted from present-day conditions to represent increased and sustained forest vulnerability — for instance, under extended droughts or other climate-driven stressors. In the model, this corresponds to increasing the average time that trees remain in the vulnerable state. Under such conditions, the system becomes more reactive and transitions into a new regime dominated by regularly recurring outbreaks. In this regime, the endemic equilibrium becomes either inaccessible or unstable, and the system exhibits recurrent boom-and-bust dynamics, which arise dynamically from the underlying feedbacks among beetle population growth, host depletion, and forest recovery. Altogether, our findings suggest that forest systems under persistent stress might become increasingly prone to self-sustained outbreak dynamics, a scenario consistent with projected impacts of climate change, foreseeing expected increases in both frequency and magnitude of extreme events [10]. Secondary spruce forests, often under suboptimal climatic conditions, have been suggested to be unsustainable in the long term due to climate warming [24].

We adopt the recurrent outbreak regime as a future scenario to investigate how external environmental shocks and management might affect or mitigate outbreak dynamics. Environmental disturbances are modeled as the conversion of a random fraction of healthy trees into susceptible ones at randomly occurring times, representing punctuated events such as storms and droughts. More frequent and intense disturbances lead to smaller but more frequent ESBB outbreaks, consistent with empirical observations of increased beetle activity during epidemic phases [44] and with evidence that repeated stressors increase forest susceptibility [26, 38]. This pattern aligns with recent recurrent outbreaks driven by rising temperatures and drought, which enhance the conversion of healthy trees into susceptible ones. In Central Europe, the combination of recurrent extreme events has increased overall forest vulnerability, leading to a prolonged, continental-scale outbreak [20, 45]. Overall, this analysis highlights the effect of the interplay between external repeated environmental stressors and internal forest dynamics.

Scheduled forest maintenance is modeled as the discrete removal of a fixed fraction of trees at regular intervals of time. Our results indicate that, in the model-predicted regime of recurrent outbreaks under increased ecological stress, management effectiveness depends not only on the removal intensity but also on timing: outbreaks can be reliably suppressed when the management removal frequency matches the system’s natural recurrence timescale. Moderately intensive but well-timed management can thus outperform frequent, low-intensity interventions. This timing sensitivity, rooted in the model’s dynamical nature, captures the interplay between internal ecological feedbacks and external interventions. Our insights complement findings from more detailed models [46], providing a mechanistic understanding of forest–management interactions and emphasizing the importance of timing-aware strategies.

Future steps include calibrating and testing the model against empirical outbreak data to infer site-specific parameters and improve predictive accuracy. Such applications require long-term data, which are not yet widely available but are increasingly being collected (D. Nardi, pers. comm.). Moreover, the framework could be extended by incorporating spatial structure — for example, through explicit modeling of beetle movement and forest heterogeneity — to investigate outbreak propagation and the effectiveness of spatially targeted interventions. Integrating spatial dynamics would enable the exploitation of geophysical and remote-sensing data, and help to predict spatial patterns of ESBB outbreaks under different environmental scenarios [47–49]. This could support the design of spatially targeted management strategies, such as optimizing trap tree deployment, evaluating passive management strategies, or exploring how mixed-species stands mitigate outbreak severity under changing disturbance regimes [50, 51].

## ACKNOWLEDGMENTS

A.D., S.A. and A.M. were supported by the Italian Ministry of University and Research (project funded under the National Recovery and Resilience Plan (NRRP), Mission 4, Component 2 Investment 1.4 - Call for tender No. 3138 of 16 December 2021, rectified by Decree n.3175 of 18 December 2021 of Italian Ministry of University and Research funded by the European Union – NextGenerationEU; Award Number: Project code CN_00000033, Concession Decree No. 1034 of 17 June 2022 adopted by the Italian Ministry of University and Research, CUP C93C22002810006, Project title “National Biodiversity Future Center - NBFC”). D.Z. gratefully acknowledges MUR and EU-FSE for financial support of the PhD fellowship PON Research and Innovation 2014-2020 (D.M 1061/2021) XXXVII Cycle / Action IV.5 “Tematiche Green”. D.N. was supported by the European Union’s Horizon Europe research and innovation programme under grant agreement No. 101134200 (FORSAID). Andrea Battisti was supported by Regione del Veneto DGR 1691-29/11/2021. The authors thank Massimo Faccoli and Giacomo Barzon for their helpful correspondence and insightful discussions.

## DATA AVAILABILITY

All data and code supporting the results of this study are available https://doi.org/10.5281/zenodo.17432288.

