## Supplementary material for "A minimal model of Norway spruce - bark beetle outbreak dynamics"

### Supplementary Information for *A minimal model of Norway spruce–bark beetle outbreak dynamics*

(Dated: October 24, 2025)

[†] These authors also contributed equally to this work.

#### SI I. FROM INDIVIDUAL-LEVEL PROCESSES TO SYSTEM-LEVEL RATES

We define the model parameters starting from a set of basic ecological processes, represented as effective reactions between individual agents. These include tree recruitment, natural weakening and death, as well as beetle-driven attacks and reproduction. The individual-level processes are listed in Table SI I, where $B$ denotes a reproductive female beetle, $T$ a healthy tree (not susceptible to colonization under endemic conditions), and $W$ a vulnerable tree. The $W$ compartment includes structurally compromised or fallen individuals typically attacked during endemic phases, as well as trees that have undergone mass beetle attacks and become vulnerable to colonization. The symbols $\varnothing$ and 0 represent, respectively, an unoccupied vegetation site and an empty site in the beetle compartment. We assume a finite carrying capacity for the tree population, such that $T + W + \varnothing = M$, where $M$ is the maximum number of tree sites in the forest patch. In contrast, we do not impose a saturation limit on the beetle population. Our model assumes a well-mixed forest patch, i.e. individuals are assumed to be homogeneously distributed across the patch, implying that the probability of interaction does not depend on spatial location. This is a common simplifying assumption in compartmental ecological models [1, 2], allowing to focus primarily on the temporal dynamics. Because insects are thus uniformly distributed across the M tree sites, it is natural to define the densities with respect to the number of tree sites: $x = \#B/M$, $y = \#T/M$, $z = \#W/M$. A schematic of the model compartments and transitions is presented in Fig. SI I.

| | |
|---|---|
| $T(W) + \varnothing \xrightarrow{\tilde{\eta}} T + T$ | birth of a healthy tree from seed dispersal of both healthy and vulnerable trees |
| $T \xrightarrow{\epsilon} W$ | healthy tree spontaneously becomes vulnerable |
| $W \xrightarrow{\rho} 0$ | vulnerable tree death |
| $B \xrightarrow{\delta} 0$ | bark beetle death |
| $B + T \xrightarrow{\tilde{\kappa}\Theta(\#B-\tilde{\sigma})} B + W$ | bark beetle attacks and weakens a healthy tree (mass attack) |
| $B + W \xrightarrow{\tilde{\beta}} nB + \varnothing$ | bark beetle colonizes a susceptible tree and reproduces, killing it |
| $0 \xrightarrow{\lambda} B$ | bark beetle immigration |

TABLE S1. Reaction rules for the Norway spruce–bark beetle model.

From the individual-level dynamics, we derive the deterministic dynamical equations in the population densities as the leading order of the Kramers–Moyal expansion of the master equation [3]:

$$\begin{cases} \dot{x} = \lambda - \delta x + \beta x z \\ \dot{y} = -\epsilon y - \kappa \Theta(x-\sigma) x y + \eta(1-y-z)(y+z) \\ \dot{z} = \epsilon y - \rho z + \kappa \Theta(x-\sigma) x y - \gamma x z \end{cases} \tag{S1}$$

The parameters $\tilde{\eta}$, $\tilde{\kappa}$, $\tilde{\sigma}$, and $\tilde{\beta}$ refer to rates or thresholds defined at the individual-tree level. To express the system in terms of normalized densities, they are rescaled as follows: $\beta = nM\tilde{\beta}$, $\gamma = M\tilde{\beta}$, $\kappa = \tilde{\kappa}$, $\eta = \tilde{\eta}M$, $\sigma = \tilde{\sigma}/M$. Notice that $\gamma$ and $\beta$ need not be equal, depending on the choice of scales of the model: indeed, if the dynamical variables signify individual densities, it will hold $\beta \gg \gamma$. Empirical estimates for these parameters are discussed in SI II.

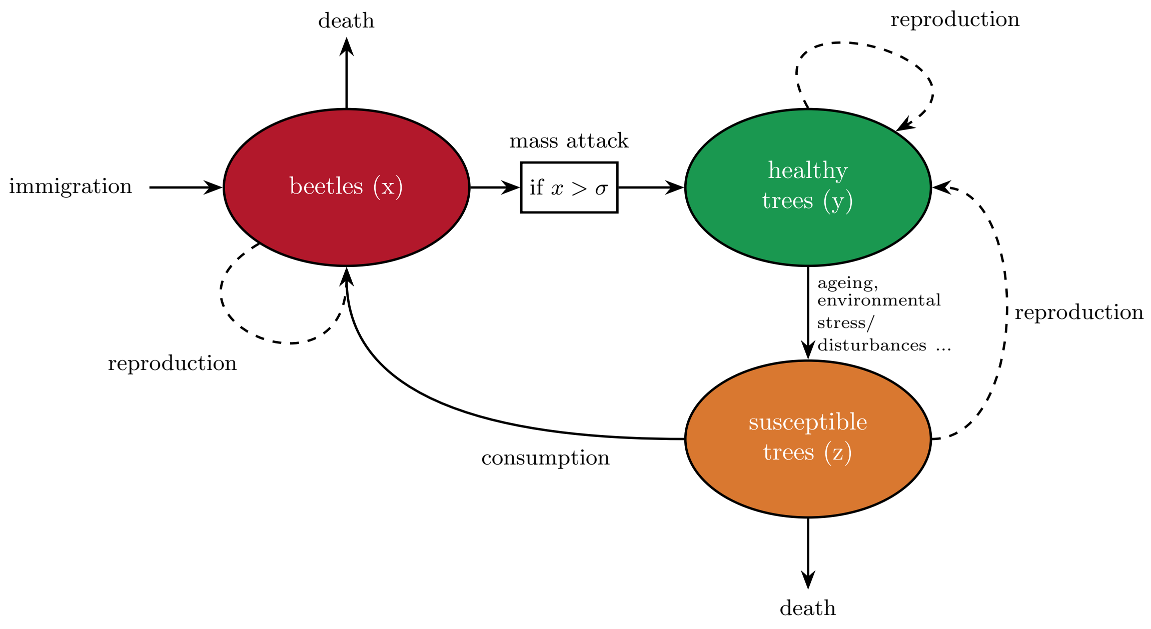


FIG. S1. **Schematic representation** of the compartmental model describing interactions between spruce bark beetles (ESBB) and host trees. Solid arrows represent transitions between states: immigration and mortality for beetles ($x$), transitions from healthy ($y$) to susceptible ($z$) trees due to aging or stress, ESBB-induced mass attacks above a density threshold $\sigma$, and mortality of susceptible trees. The reproductive role of each compartment is shown with dashed arrows. Vulnerable trees serve as the primary resource for beetle reproduction and are depleted through consumption.

#### SI II. REFERENCE PARAMETERS ESTIMATION

To calibrate the model, we estimate reference values for the parameters based on available ecological and entomological literature. These estimates reflect typical conditions in managed European spruce forests, while also accounting for known sources of variability. Where precise data are lacking, we use informed assumptions to define plausible orders of magnitude or leave parameters free for exploration. Table I of the main text summarizes the final reference values adopted in the model, along with estimated variability ranges. These ranges are derived from ecological heterogeneity (e.g., altitude, forest density) or uncertainty in empirical measurements.

Several simplifying assumptions have been made for tractability. First, although bark beetles are largely inactive during winter, we model processes as if they occur continuously throughout the year, effectively averaging over seasonal dynamics. Second, we assume two generations per year, reflecting average life cycle patterns observed across altitudes and typical annual temperatures. Third, since only female beetles contribute to reproduction, we track uniquely female individuals; consequently, parameters involving reproduction (notably $\beta$) are adjusted by assuming a 1:1 sex ratio and halving the brood size accordingly. Tree recruitment, modeled by a logistic curve, provides a mechanism for host replenishment and stand saturation effect. Healthy trees are assumed to transition through a vulnerable state before dying, thus the typical lifetime of an individual tree, $t_{life}$, can be approximated as the sum of the average durations spent in each compartment, i.e., $t_{life} \approx \epsilon^{-1} + \rho^{-1}$.

##### Tree demographic parameters

*Maximum number of trees per patch (M)*

We normalize the system to a forest patch of 1 hectare, assuming a typical total number of $M \sim 500$ spruce trees per patch [4]. Observed values usually vary from 300 in high-altitude or sparse stands to 700 in dense, low-altitude forests (G. Petit, pers. comm.). Where relevant, we explicitly relate individual-level processes (e.g., fecundity) to system-level rates via rescaling to a forest patch of area 1 ha, as explained in SI I. This should not be taken to mean that our model represents a forest patch of 1 ha: as we are making the assumption of spatial homogeneity, the results can in principle be scaled up to a forest in any size. Indeed, it is rather the assumption of homogeneity that restricts our model by neglecting spatial dynamics at all scales.

*Death rate of susceptible trees ($\rho$)*

Susceptible trees represent vulnerable individuals that include both structurally compromised trees, such as those heavily damaged or uprooted by wind (but we may include senescent, severely drought-stressed or pathogen-stressed trees as well) and otherwise healthy trees that have undergone mass beetle attacks. Wind-fell trees are typically expected to degrade and die mostly within a few years [5]. Based on this, we assume an average viability of about 1 to 4 years following the onset of weakening, yielding a plausible range of $6 \cdot 10^{-4}$ to $5 \cdot 10^{-3}$ day$^{-1}$, which reflects variability in environmental conditions.

*Conversion rate from healthy to susceptible trees ($\epsilon$)*

Assuming that trees typically pass through a vulnerable state before dying, we estimate $\epsilon$ based on typical lifespans. For a typical lifespan of 250 years, within a conservative range of 100–400 years [6, 7], this yields $\epsilon = 1 \cdot 10^{-5}$ day$^{-1}$. The plausible range from $7 \cdot 10^{-6}$ to $3 \cdot 10^{-5}$ day$^{-1}$ reflects such variation in tree longevity and site-specific factors.

*Birth rate of healthy trees ($\eta$)*

Tree recruitment rates depend on forest age structure and site productivity. We assume that the growth of the tree population can be modeled as a logistic curve, with growth rate $\tilde{\eta} = 1/(100 \text{ years})$, and scale by $M$ to obtain $\eta = 1.3 \cdot 10^{-2}$ day$^{-1}$. This parameter reflects the rate at which new trees reach a proper size such that they can be attacked by ESBB. Based on regeneration times ranging from about 60 years in low-altitude stands to over 200 years in high-altitude forests (G. Petit, pers. comm.), we define a plausible range between $4 \cdot 10^{-3}$ and $3 \cdot 10^{-2}$ day$^{-1}$.

**Beetle demographic parameters**

*Death rate of beetles ($\delta$)*

We assume a typical adult flight of 1–3 days during the dispersal phase, leading to $\delta = 1/(2 \text{ days}) \approx 0.5$ as the reference value. This corresponds to a plausible range between between 0.3 and 1.0 day$^{-1}$.

*Beetles per brood ($n$)*

Each female typically lays around 80 eggs [8]. Since only a fraction of individuals survive the winter and successfully emerge as adults, we account for an estimated winter mortality of 30% to 50% [8] (D. Nardi, pers. comm. from field observations). Additionally, assuming a 1:1 sex ratio, we consider only the female offspring contributing to reproduction, effectively dividing the brood size by two. This yields an average of $n \approx 25$ viable female offspring. We allow for a range from 20 to 30 to reflect variability in survival due to environmental conditions.

*Reproduction rate of beetles ($\beta$)*

We assume two generations per year. As detailed in SI II B, we compute the growth rate by matching the population growth resulting from two successive cohorts in a single year with a continuous time growth, obtaining $\beta \approx 18$ day$^{-1}$. We consider a range from 5 to 40 day$^{-1}$ to account for variation in brood size, survival, and generation time. We remark that here we assumed conditions which are favorable to the growth of beetle population, neglecting interspecific competition.

**Beetle-tree interaction parameters**

*Tree death rate due to beetle attack ($\gamma$)*

This rate is linked to $\beta$ via $\gamma = \beta/n$. Given variability in $\beta$ and $n$, we define a range of0.1 to 0.7 day$^{-1}$, with a reference value of 0.4 day$^{-1}$.

*Threshold for mass attack ($\sigma$)*

Mass attacks on healthy trees require a critical number of insects, typically estimated at around $\tilde{\sigma} \sim 5000$ individuals per tree [9]. In the model, this threshold is expressed as a condition on beetle density, using a scaling argument that accounts for local aggregation behavior. Due to pheromone-mediated chemical signaling, ESBB are known to aggregate over a finite spatial scale. Lacking data on natural aggregation distances, we use artificial chemical signaling [10] as a broad estimate of aggregation radius; where the effective pheromone attraction range is estimated to lie in the range $R \sim 19$–$97$ m. As a result, each tree is not exposed to the entire beetle population in the forest, but only to those within an aggregation area $A \sim \pi R^2$ out of the total area $L^2$. Accordingly, we assume that mass attacks are triggered when the number of insects within this aggregation area exceeds $\tilde{\sigma}$. This leads to an effective density threshold $\sigma = \tilde{\sigma}/m$, where $m$ is the number of trees within the aggregation area: $m = M \cdot A/L^2$. Based on empirical attraction ranges, we estimate that $\sigma$ lies between approximately 3.3 and 83. Thus, given this uncertainty, we set as a conservative reference value $\sigma = 10$. A detailed derivation of this scaling and its implications is provided in SI II A.

*Colonization rate of healthy trees ($\kappa$)*

This parameter reflects the efficiency with which ESBB successfully colonizes and weakens healthy trees once mass attacks are triggered. This is distinct from the threshold parameter $\sigma$, which controls the number of insects required to initiate an attack. Both $\kappa$ and $\sigma$ can be modified to reflect the health state of the forest, drought, and prior damage: while $\sigma$ captures the resistance threshold of trees, $\kappa$ describes how effectively insects exploit that vulnerability under outbreak conditions. Due to the lack of direct empirical estimates, we treat $\kappa$ as a free parameter in our scenario analysis.

##### A. Density threshold for mass attacks and scalability of the model

Mass attacks on healthy trees typically require around $\tilde{\sigma} \sim 5000$ beetles per tree. In the model, this threshold must be expressed in terms of beetle density $x = B/M$. Different assumptions about aggregation lead to different scalings of the Heaviside function used to trigger mass attacks.

*A) Global aggregation:* we assume that all beetles in the forest patch can potentially attack a single tree. Then:

- $\#B =$ total number of beetles in the forest patch,
- $\tilde{\sigma} =$ number of beetles needed to colonize a healthy tree,

leading to the condition: $\Theta(\#B - \tilde{\sigma}) \propto \Theta\left(x - \frac{\tilde{\sigma}}{M}\right)$.

*B) Local density threshold:* we assume that each tree is only affected by the local beetle density in its immediate vicinity:

- $x = \#B/M =$ average number of beetles per tree site,
- $\tilde{\sigma} =$ beetle threshold per tree,

resulting in the simpler form: $\Theta\left(x - \tilde{\sigma}\right)$. This implies beetles do not aggregate and each tree interacts with the local beetle density only.

*C) Intermediate aggregation scale:* a more realistic scenario accounts for beetle aggregation over a finite spatial range. Beetles are known to attract mates via pheromone-mediated chemical signaling over a certain spatial length-scale, which we denote by $R$. This leads to an effective aggregation area $A = \pi R^2$, which defines a fraction $f = A/L^2$ of the total forest patch corresponding to $m = f \cdot M = AM/L^2$ tree sites. Hence, we compare:

- $x\, m = \#B\, m/M =$ average number of beetles within the aggregation area,

- $\tilde{\sigma}$ = per-tree attack threshold,

yielding: $\Theta(xm - \tilde{\sigma}) \propto \Theta\left(x - \frac{\tilde{\sigma}}{m}\right)$. Since $m$ is determined by the aggregation area $A$ and not the total patch size $L^2$, this formulation captures beetle spatial behavior while preserving scalability with forest patch size.

##### B. Continuous-time consistency in determining $\beta$ and $\gamma$

We wish to estimate $\beta$ from empirical observations, and consequently derive $\gamma$. However, a considerable difficulty is given by the fact that $\beta$ is not merely a birth rate for ESBBs; instead, the dynamical birth rate at time $t$ is given by $\beta z(t)$, which depends on the availability of susceptible trees. As such, empirical estimates on biological grounds are not readily available, and we must root our analysis in population dynamics.

Our starting point is the observation that each female ESBB entering a susceptible tree reproduces, yielding a net increase in ESBB population. Furthermore, in each year, a succession of $N$ cohorts will be observed, with $1 \leq N \leq 3$. In the following we will neglect all other processes, and work under the assumption that, through successive reproductive cycles, each offspring will reproduce with similar success. Under these assumptions, if $N$ cohorts emerge in a time $T$, with each individual generating $n$ offspring, we expect that during this time the population of ESBB will increase by a factor $\mu \approx n^N$. Hence, by tracking the growth rate of successive cohorts and matching the resulting population growth, we are able to link expected populations levels to likely values of $\beta$.

For the sake of conceptual clarity, we first estimate $\beta$ by assuming the continuous time evolution for ESBB density, and assuming that the density of susceptible trees does not change over time, namely:

$$\dot{x} = \beta x z, \quad z = \text{ const.} \tag{S2}$$

In the time interval $[0, T]$, the population of ESBB increases of a factor $\mu$, where

$$\mu = \frac{x(T)}{x(0)} = e^{\beta z T} \tag{S3}$$

Thus, knowing the parameters $\mu$ and $T$, we can estimate

$$\beta = \frac{\log(\mu)}{z\,T}. \tag{S4}$$

The use of estimate (S4) in a continuous time model leads to the requested growth of ESBB population over a given time $T$. However, this estimate is obtained under the assumption that $z$ is constant, it is not clear what value of $z$ one should utilize; yet, this value will directly affect our estimation of $\beta$. Both these difficulties are addressed by taking into account the dynamic nature of $z$, as obtained by the following differential equations:

$$\begin{cases} \dot{x} = \beta x z \\ \dot{z} = -\gamma x z \quad \text{with } \gamma = \beta/n \end{cases} \tag{S5}$$

Through eqs. (S5), we manifestly tie together the dependence of $\beta$ on $n$ and the non-reciprocity of $x - z$ interaction; however, we note that this does not reduce the number of independent parameters. The ESBB relative growth, $x(T)/x(0) = \mu$, is easily obtained as solution of eqs. (S5) by noting that $C = x + nz$ is a conserved quantity:

$$\mu = x(T)/x(0) = \frac{C/x(0)}{1 + \frac{nz(0)}{x(0)} e^{-(\beta C/n)T}} \,. \tag{S6}$$

From this, $\beta$ can be extracted in terms of $x(0)$, $z(0)$, $T$ $n$ and $\mu$: we find

$$\beta = \frac{n}{CT} \log\left(\frac{C/x_0 - 1}{C/(x_0\mu) - 1}\right) \,. \tag{S7}$$

In the limit of $x(0) \to 0$, we recover the result in Eq. (S4) (which is independent of $n$), provided we identify $z \equiv z(0)$. Here, the assumption of constant density of vulnerable trees is substituted with the assumption of small initial ESBB density, consistent with an endemic state. Therefore, $z(0)$ should reflect a low density of susceptible trees in standard conditions, namely (see Table I) $z(0) = \frac{\text{few susceptible trees}}{L} / \frac{M \text{ trees}}{L} \propto M^{-1}$. As it is impossible — and perhaps not meaningful — to assign a fixed value to $z(0)$, we will consider a rather broad range of values for $\beta$ comprising the estimates obtained in this section.

#### SI III. EQUILIBRIUM STATES

Mathematically, equilibria (or *fixed points*) are defined as the solutions of the dynamical equations such that their time derivatives vanish [1]. To determine the equilibria of the system, therefore, we have to calculate the solutions of the equations Eq. (S1) where the left-hand side is equal to zero. We denote an equilibrium as $(x^*, y^*, z^*)$, corresponding to the stable densities of ESBB, vulnerable trees, and healthy trees, respectively. We classify them according to whether the ESBB abundance lies below or above the outbreak threshold $\sigma$.

In the sub-threshold regime ($x^* < \sigma$), two possible equilibria exist.

1. The trivial, ecologically unrealistic equilibrium is:

$$(x^*, y^*, z^*) = (\delta/\lambda, 0, 0) \; . \tag{S8}$$

2. The endemic coexistence equilibrium, where beetles and trees persist, is found by solving the following equation in the beetle population $x^*$:

$$f_<(x^*) = 0, \qquad f_<(x) = a_3 x^3 + a_2 x^2 + a_1 x + a_0. \tag{S9}$$

where $f_<(x)$ is a 3-rd order polynomial in the ESBB density $x$, whose coefficients depend on the model's ecological parameters as follows:

$$\begin{aligned}
a_3 &= \gamma^2 \, \delta \, \eta \\
a_2 &= \gamma \left[\beta\epsilon(\epsilon - \eta) - \gamma \, \eta \, \lambda + 2\delta\eta(\rho + \epsilon)\right] \\
a_1 &= \eta(\rho + \epsilon)(\delta(\rho + \epsilon) - 2\gamma\lambda) - \beta\epsilon(\eta(\rho + \epsilon) - \rho\epsilon) \\
a_0 &= -\eta\lambda(\rho + \epsilon)^2
\end{aligned} \tag{S10}$$

The corresponding tree densities of this endemic equilibrium are given by:

$$z^* = \frac{\delta x^* - \lambda}{\beta x^*}, \quad y^* = z^* \, \frac{\rho + \gamma x^*}{\epsilon} \; . \tag{S11}$$

Above the outbreak threshold ($x^* > \sigma$), mass attacks on trees alter the system dynamics, leading to a modified fixed point. ESBB equilibrium abundance is given by the solution $x^*$

of $f_>(x^*) = 0$, where $f_>(x)$ is a quartic polynomial incorporating the additional effects of beetle-induced tree mortality through the attack rate $\kappa$:

$$f_>(x^*) = 0, \qquad f_>(x) = b_4 x^4 + b_3 x^3 + b_2 x^2 + b_1 x + b_0. \tag{S12}$$

with coefficients:

$$\begin{aligned}
b_4 &= \beta\gamma\kappa^2 \\
b_3 &= \gamma\kappa(-\beta\eta + 2\beta\epsilon + 2\delta\eta) + \kappa^2(\beta(\rho - \eta) + \delta\eta) + \gamma^2\delta\eta \\
b_2 &= \beta\epsilon(\gamma(\epsilon - \eta) - 2\eta\kappa) + \beta\kappa\rho(2\epsilon - \eta) - \eta(\gamma + \kappa)(\lambda(\gamma + \kappa) - 2\delta(\rho + \epsilon)) \\
b_1 &= \eta(\rho + \epsilon)(\delta(\rho + \epsilon) - 2\lambda(\gamma + \kappa)) - \beta\epsilon(\eta(\rho + \epsilon) - \rho\epsilon) \\
b_0 &= -\eta\lambda(\rho + \epsilon)^2
\end{aligned}$$

In this regime, tree densities are correspondingly given by:

$$z^* = \frac{\delta x^* - \lambda}{\beta x^*}, \quad y^* = z^* \frac{\rho + \gamma x^*}{\kappa x^* + \epsilon} \tag{S13}$$

Given the full expression of the coefficients, Eq. (S9) and Eq. (S12) can be solved numerically, yielding the fixed points $(x^*, y^*, z^*)$. To analyze their stability, we compute the Jacobian matrix $\mathbb{J}(x^*, y^*, z^*)$, which describes the local linearization of the system around equilibrium points. The explicit expression for $\mathbb{J}(x, y, z)$ is given by:

$$\mathbb{J}(x,y,z) = \begin{pmatrix} \beta z - \delta & 0 & \beta x \\ -\kappa y\left[x\delta(x-\sigma) + \theta(x-\sigma)\right] & \eta\left[1 - 2(y+z)\right] - \kappa x\theta(x-\sigma) - \epsilon & \eta\left[1 - 2(y+z)\right] \\ -\gamma z + \kappa y\left[x\delta(x-\sigma) + \theta(x-\sigma)\right] & \epsilon + \kappa x\theta(x-\sigma) & -(\gamma x + \rho) \end{pmatrix} \tag{S14}$$

where $\delta(x - \sigma)$ denotes the Dirac delta function, and $\theta(x - \sigma)$ is the Heaviside step function. The eigenvalues of $\mathbb{J}(x, y, z)$ determine the local stability properties of each fixed point. While we have obtained analytical expressions for the eigenvalues, their complexity makes them impractical to present explicitly. There are three possible scenarios. In the first scenario there is at least one eigenvalue with positive real part, which corresponds to an unstable equilibrium. In the second scenario all the eigenvalues are real and negative, which corresponds to a stable equilibrium that is reached monotonically exponentially at large times. In the third scenario all eigenvalues have negative real part, but a couple of them have nonzero imaginary part. The latter scenario corresponds to a stable equilibrium that is reached with a damped oscillation at large times.

#### SI IV. REACTIVITY

Reactivity assesses a system's transient response to perturbations. It is determined by the largest eigenvalue of the symmetric part of the Jacobian matrix:

$$\mathbb{J}_{\text{symm}} = \frac{1}{2}(\mathbb{J} + \mathbb{J}^T) = \tag{S15}$$
$$= \frac{1}{2}\begin{pmatrix} 2(\beta z - \delta) & -\kappa y[x\delta(s-x) + \theta(x-s)] & \beta x - \gamma z + \kappa y[x\delta(s-x) + \theta(x-s)] \\ -\kappa y[x\delta(s-x) + \theta(x-s)] & -2[\epsilon + 2\eta(y+z-1) + \kappa x\theta(x-s)] & \epsilon - 2\eta(y+z-1) + \kappa x\theta(x-s) \\ \beta x - \gamma z + \kappa y[x\delta(s-x) + \theta(x-s)] & \epsilon + \eta[1 - 2(y+z)] + \kappa x\theta(x-s) & -2(\gamma x + \rho) \end{pmatrix}$$

While we have obtained analytical expressions for the eigenvalues of $\mathbb{J}_{\text{symm}}$, they are too complex to be presented explicitly. Instead, we focus on numerical evaluations to analyze their dependence on key parameters and to identify conditions under which transient growth is maximized.

We proceed to analyze the reactivity of the endemic coexistence equilibrium, which represents the most ecologically plausible scenario under typical conditions. We start by evaluating Eq. (S15) at the reference parameter values reported in Table I of the main text. The eigenvalues of $\mathbb{J}_{\text{symm}}$ at the reference endemic equilibrium are approximately

$$\lambda_1 \sim 0.002,\ \lambda_2 \sim -0.02,\ \text{and}\ \lambda_3 \sim -0.4\ . \tag{S16}$$

The positive value of $\lambda_1$ indicates that such equilibrium is reactive. To illustrate the reactive behavior, we numerically integrate the system after applying small perturbations (of magnitude $\epsilon = 0.05$) along each eigenvector of $\mathbb{J}_{\text{symm}}$. The most reactive direction (associated with $\lambda_1$ ) leads to a transient spike in the perturbation norm $||\delta r(t)||$, while non-reactive directions, i.e. all the ones perpendicular to the reactive direction, result in monotonic relaxation (Fig. 2a of the main text). The dominant eigenvector is approximately

$$
\mathbf{v}_1 \sim \begin{bmatrix} 0.05 \\ -0.40 \\ 0.91 \end{bmatrix}, \tag{S17}
$$

indicating that transient growth is primarily driven by shifts in the weakened tree population. In contrast, directions in the plane spanned by

$$
\mathbf{v}_2 = \begin{bmatrix} 0.02 \\ 0.9 \\ 0.4 \end{bmatrix} \quad \text{and} \quad \mathbf{v}_3 = \begin{bmatrix} -0.9987 \\ 0.0008 \\ 0.0504 \end{bmatrix} \tag{S18}
$$

are dominated by changes in healthy trees or beetles alone and do not produce significant transient growth. This indicates that reactivity in the endemic equilibrium is mainly driven by perturbations involving the abundance of vulnerable trees. To further illustrate ecological relevance, Fig. 2b of the main text shows the full system trajectory following a perturbation with a strong projection along $\mathbf{v}_1$, (with $\epsilon = 0.1$).

We report below additional reactivity maps visualizing reactivity's dependence on ecological parameters.

#### SI V. RECURRENT OUTBREAKS

To study the recurrent outbreak regime, we choose parameter values that give rise to this particular dynamical behavior. These values are outside the ecologically plausible range representing current forest conditions (Table I of the main text), but we focus on the case in which they still preserve a realistic balance between key ecological rates. In the selected parameters case, an equilibrium exists but it is only weakly stable due to its high reactivity, making it effectively inaccessible without fine-tuning of the initial conditions: even minimal increases in the number of susceptible trees can push the system away from equilibrium and trigger outbreaks.

Because the insect population periodically collapses, allowing susceptible trees to accumulate during intervals when it is effectively absent, we consider a regime of weak immigration by setting $\lambda \gtrsim 0$. After recurrent outbreaks begin, $\lambda$ plays no further role. Here, the stability properties of the system are effectively unaltered with respect to the case $\lambda = 0$. Thus,

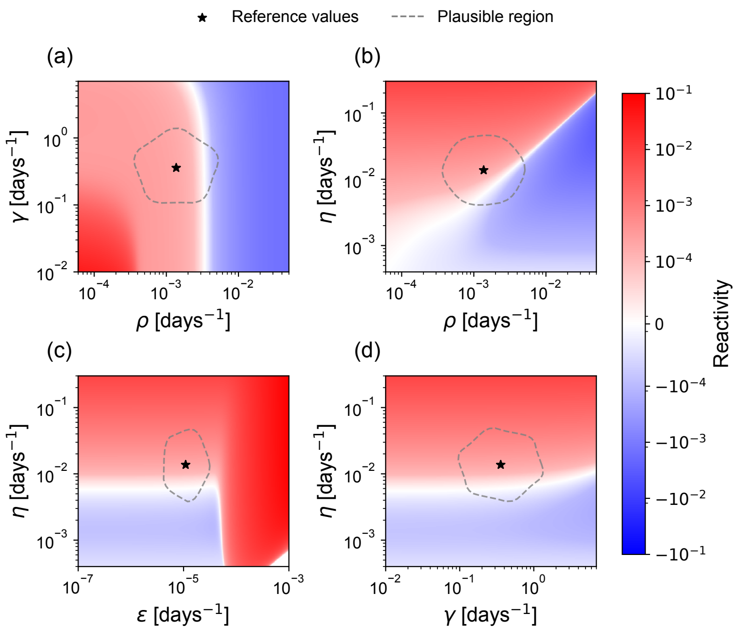


FIG. S2. Reactivity landscape over a region of parameter space centered around the reference values given in Table I of the main text. The star indicates the reference parameter set, for which the reactivity is $R \approx 0.002$. Within the ecologically plausible region, reactivity is predominantly positive. Therefore, despite parameter values and ranges being approximate, the system is generally expected to exhibit reactive dynamics under realistic ecological conditions, indicating an inherent vulnerability to transient outbreaks triggered by perturbations.

we will start from initial conditions for trees' population in proximity of the ESBB-free stationary values, obtained from Eq. (S1) for $\lambda = 0 = x^*$, finding

$$x(0) = x^* = 0 \ , \quad y(0) = y^* = \frac{\rho\big[\epsilon(1-\frac{\rho}{\eta})+\rho\big]}{(\epsilon+\rho)^2} \ , \quad z(0) = z^* = \frac{\epsilon}{\rho} y(0) \ . \tag{S19}$$

This fixed point is feasible, i.e. $y^* > 0$, if $\eta > (\epsilon^{-1} + \rho^{-1})^{-1}$. Moreover, the insect population can successfully invade, i.e. has positive growth at low density, if the following condition is satisfied:

$$\beta > \frac{\delta\eta(\epsilon+\rho)^2}{\epsilon\big[\epsilon(\eta-\rho)+\eta\rho\big]} \ . \tag{S20}$$

Notably, the denominator in Eq. (S20) is proportional to $y^*$ and $z^*$, implying that invasion becomes increasingly difficult as the equilibrium number of host trees declines. Moreover, even when condition (S20) holds, it does not guarantee the emergence of recurrent outbreaks. These require additional dynamical conditions — in particular, sufficiently high reactivity

— as pointed out in Section III C of the main text.

The observables used to characterize the recurrent outbreak regime are defined as the maximum population size during an outbreak $x_{\text{peak}}$ (BOS), and the time between successive outbreaks $T$ (BWT). In such regime, the dynamics is regularly repeating, therefore they are linked by the periodicity condition :

$$x(t) = x(t+T) \equiv x_{\text{peak}} \tag{S21}$$

We underline that in this section we analyze stressors as encoded by static parameters, hence the exact periodic dynamics are to be intended as describing the overall susceptibility of the system to ESBB outbreaks, in a mean sense. Stressor anomalies, described by dynamical noise, are considered in the next section.

#### SI VI. ENVIRONMENTAL DISTURBANCES

Each disturbance is incorporated in our model as an external force that occurs at random times and with random magnitude, and converts a fraction of healthy trees into susceptible trees. A *temporal series of disturbances* is described by the sequence of random pairs

$$\left\{\left(\Delta_{(n)}, \tau_{(n)}\right)\right\}_n \ . \tag{S22}$$

Here, $\tau_{(n)}$ denotes the time of the n-th disturbance, and $\Delta_{(n)} \in (0,1)$ its magnitude — i.e., the fraction of healthy trees converted into susceptible ones. Each $\Delta_{(n)}$ is independently drawn from a probability distribution with support in $(0,1)$, whose shape is controlled by the parameter $\Delta_d$. The occurrence times $\tau_{(n)}$ are defined recursively as $\tau_{(n)} = \tau'_n + \sum_{m=0}^{n-1} \tau_{(m)}$, where each waiting time $\tau'_{(n)}$ is independently and identically sampled from an exponential distribution, $\tau'_{(n)} \sim 1/\tau e^{-\tau'_{(n)}/\tau_d}$. The parameter $\tau_d$ thus defines the (inverse) typical frequency of disturbances. For simplicity, we do not consider the general case of statistically dependent $\Delta_{(n)}$ and $\tau_{(n)}$, although this possibility could better characterize realistic disturbance [11, 12]. Nonetheless, allow us to explore how shifts in environmental conditions, for instance, due to climate change or ecological invasions, affect outbreak dynamics. Specifically, decreasing $\tau_d$ leads to more frequent disturbances, while increasing $\Delta_d$ corresponds to more severe ones. The disturbance series (S22) acts on the dynamics (S1) through the multiplicative pulse function

$$\mathcal{D}_{\Delta_d,\tau_d}(t) \equiv \sum_{n=0}^{+\infty} \Delta_{(n)} \delta\left(t - \tau_{(n)}\right) \ , \tag{S23}$$

and $\delta(\cdot)$ is Dirac's Delta function. Specifically, eqs. (S1) are extended to comprise the time-dependent terms $\dot{y}(t) = \cdots - \mathcal{D}_{\Delta_d,\tau_d}(t)y(t)$ and $\dot{z}(t) = \cdots + \mathcal{D}_{\Delta_d,\tau_d}(t)y(t)$. For mathematical consistency, we require that as $\tau_d \to 0$, $\Delta_d \to 0$ as well [13]. Under this stochastic forcing, the system no longer follows strictly periodic dynamics as in the unperturbed case: both $x_{\text{peak}}$ and $T$ become random variables whose distributions depend on the disturbance parameters. The impact of the disturbance term on tree populations can be verified directly. For instance, the term$\dot{y}(t) = \cdots - \Delta_{(n)}\delta(y - \tau_{(n)})y(t)$ yields $y(t_+) = \int_0^{t_+} dt'\, \dot{y}(t') = y(t_-)\left(1 - \Delta_{(n)}\right)$ where $t_-$ and $t_+$ are times immediately before and after the disturbance. Analogously, $z(t_+) = z(t_-) + y(t_-)\Delta_{(n)}$. As expected, each disturbance reduces healthy trees and increases susceptible ones. Intuitively, more frequent or severe disturbances are expected to reduce the average waiting time $T$ with respect to the baseline (BOWT). However, the effect on the outbreak size $x_{\text{peak}}$ is less obvious. As shown in Figure 5 (top) of the main text, the distribution of $T$ is upper-bounded by the BOWT observed in unperturbed recurrent outbreak dynamics, shifting towards smaller waiting times as $\tau_d$ decreases and $\Delta_d$ increases. Simultaneously, the distribution of $x_{\text{peak}}$ becomes broader: most outbreaks become smaller than in the baseline case, though occasional extreme events may still exceed the baseline size. Taken together, these results suggest that more frequent and severe disturbances lead to smaller but more frequent outbreaks, a shift consistent with the increased overall severity of ESBB activity during epidemic phases [14].

#### SI VII. MANAGEMENT STRATEGIES

We now consider how periodic management can reduce the risk of ESBB outbreaks. We define a *management strategy* $\mathcal{M}_{\Delta_m,\tau_m}$ as a periodic forcing that instantaneously reduces the density of susceptible trees,$z \to (1 - \Delta_m)z$, at regular time intervals $\tau_m$. This is introduced into the model by modifying the equation for $\dot{z}(t)$ as:

$$\dot{z}(t) = \cdots - \mathcal{M}_{\Delta_m,\tau_m}(t)z(t) \ , \tag{S24}$$

where $0 < \Delta_m < 1$ and $\tau_m > 0$, and we defined the forcing operator

$$\mathcal{M}_{\Delta_m,\tau_m}(t) \equiv \Delta_m \sum_{n=0}^{+\infty} \delta(t - n\tau_m) \ , \tag{S25}$$

The management strategy seeks to limit ESBB population growth by reducing the density of available susceptible trees. Without considering constraints on management, if this reduction is frequent and thorough enough, i.e. if $\tau_m$ and $\Delta_m$ are respectively small and large enough, the insect density will remain below the threshold for mass attacks on healthy trees, thus preventing outbreaks altogether. However, as we will argue next, $\tau_m$ and $\Delta_m$ may not be chosen independently from each other. To formalize this, we first examine the limit $\tau_m \to 0$, corresponding to continuous management. This limit is well-defined if $\Delta_m = \bar{\Delta}\tau_m + O(\tau_m^2)$, in which case the forcing converges to a constant rate $\mathcal{M}_{\bar{\Delta}\tau_m,\tau_m}(t) \to \bar{\Delta}$. The susceptible-tree equation becomes: $\dot{z} = \cdots - (\rho + \bar{\Delta})z$ and outbreaks are avoided if the invasion condition from Eq. (S20) is violated under the substitution $\rho \to \rho + \bar{\Delta}$. From a practical standpoint, this scaling is also realistic: high-frequency interventions can only remove a small fraction of susceptible trees at a time. To account for the previous mathematical and practical considerations, we choose

$$\Delta_m = \frac{\hat{\Delta}\tau_m}{\tau_0 + \tau_m} \ , \tag{S26}$$

where $\hat{\Delta} < 1$ and the constant $\tau_0$ defines the time scale past which waiting for the next intervention has reduced marginal utility. In this case, $\Delta_m < \hat{\Delta}$ for any finite $\tau_m$, while for $\tau_m \sim 0$ we may approximate $\Delta_m = (\hat{\Delta}/\tau_0)\tau_m + O(\tau_m^2)$. Then, the previous case of linear scaling is recovered for $\tau \ll \tau_0$ with the identification $\bar{\Delta} = \hat{\Delta}/\tau_0$. While $\bar{\Delta}$ has the dimension of an inverse time, since $\tau_0$ is a time scale, $\hat{\Delta}$ is a dimensionless quantity. We now examine the effectiveness of such strategies in the presence of small and frequent disturbances (Figure 5 (bottom) of the main text). In the absence of management, outbreak observables $x_{\text{peak}}$ and $T$ remain tightly distributed around the baseline values (BOS and BOWT). In the limit $\tau_m \to 0$, where vulnerable trees are continuously removed, management is ineffective unless $\hat{\Delta}/\tau_0$ is very large. More realistically, when $\hat{\Delta}/\tau_0$ is small, outbreaks can only be prevented if $\tau_m \gtrsim$ BOWT. Beyond this threshold, the ecological response to management becomes irregular: both $x_{\text{peak}}$ and $T$ switch between non-outbreak conditions (where $T = 0$) and and their maximum value, as $\tau_m$ increases together with $\Delta_m$ (Eq. (S26)). However, we find invariably that management is reliably effective when $\tau_m$ is slightly larger than $\langle T' \rangle$, which represents the average outbreak waiting under management. Therefore, in this regime of large $\tau_m$, the success of a management strategy depends not only on the magnitude of intervention, but also on the timing which must align with the system's intrinsic response timescale.

#### SI VIII. PARAMETERS AND OTHER DETAILS ON FIGURES

**General information.** Unless specified otherwise, Eq.(1) has been integrated using a fourth-order Runge-Kutta scheme. Whenever necessary, peaks have been automatically detected using the function `scipy.signal.find_peaks`.

**Fig. 1.** Trajectories are obtained by integrating Eq.(1) using the Euler scheme. The parameter values used are reported in Table I, and the reactivity associated to the stable fixed point $(x^-, y, z)$, representing the endemic state, is $\lambda \simeq 2 \cdot 10^{-3}$.

**Fig 2.** (b) For each stable fixed point across parameter space, determined analytically as explained in Section IIC, we calculate reactivity (Section IID) as detailed in SI IV. The parameter values kept fixed in these diagrams are reported in Table I.

**Fig. 3.** The phase diagrams here presented are analytical. The fixed points displayed are obtained by studying the existence and stability of the stationary solutions presented in Section IIC and detailed in SI III. (a) The parameter values kept fixed in this diagram are: $\delta = 5$, $\beta = 10$, $\sigma = 5$, $\kappa = 1$, $\epsilon = 10^{-2}$, $\gamma = 10^{-2}$, $\rho = 10^{-2}$, $\eta = 0.5$, $\lambda = 10^{-5}$. (b) The parameter values used in these diagrams are reported in Table I. (c) Trajectories have been numerically integrated using the Euler scheme or the fourth-order Runge-Kutta scheme (in the case of oscillatory trajectories).

**Fig. 4.** For both panels (a) and (b), $\lambda = 10^{-200}$, $\delta = 0.5$, $\eta = 5 \times 10^{-3}$, $\epsilon = \rho = 10^{-5}$, $\gamma = 10^{-2}$, $\kappa = 10^{-3}$, $\sigma = 10$; $dt = 0.1$, $t_{\max} = 365 \times 100$. (a) $\beta = 20$; (b): $\beta$ varies over the interval $[5, 30]$. Initial conditions are sampled ((a): $5 \times 10^3$ samples; (b): $10^3$ samples) by perturbing the endemic fixed point found numerically (see Section IIC) with a Gaussian array with independent components of zero mean, and variance equal to $10^{-3}$. Negative components of initial conditions, whenever found, have been set equal to zero. Periodic solutions have been detected by counting peaks of $x(t)$ exceeding a given density threshold ($\sigma$ for both (a) and (b), with also smaller thresholds for (b)). (c) Parameters are as in

Fig. 4 (a-b), with fixed $\beta = 20$, whereas initial conditions are given by Eq. (S19). (d) Same parameters and initial conditions as panel (c), but with varying $\kappa \in [10^{-3}, 2 \times 10^{-2}]$.

**Fig. 5.** (top) Models' parameters are identical as in Fig. 4 (c-d); $dt = 0.05$, $t_{\max} = 365 \times 10^3$. (bottom) Figures are obtained by the simulations of Eq. (1), replicated 1000 times, with all model's parameters and initial conditions equal to those of Fig. 4 (a-b). The parameters of management $\mathcal{M}_{\Delta,\tau}$ with $\Delta = \frac{\hat{\Delta}\tau}{\tau_0+\tau}$ are $\hat{\Delta} = 1.8 \times 10^{-4}$ and $\tau_0 = 10$ years.

**Fig. S2.** The plots here presented are analytical. The fixed points are obtained by studying the existence and stability of the stationary solutions presented in Section IIC and detailed in SI III, while the associated reactivity (Section IID) is calculated as explained in SI IV. The parameter values used are reported in Table I.

---


[1] James D. Murray. *Mathematical Biology I. An Introduction*, volume 17 of *Interdisciplinary Applied Mathematics*. Springer, New York, 3 edition, 2002. doi:10.1007/b98868.

[2] P.B. Turchin. *Complex population dynamics: A theoretical/empirical synthesis*, volume 35. 01 2003. doi:10.1515/9781400847280.

[3] Nicolaas Godfried Van Kampen. *Stochastic processes in physics and chemistry*, volume 1. Elsevier, 1992.

[4] Ludwig Olofsson, Ola Langvall, and Arne Pommerening. Norway spruce (*Picea abies* (L.) H. Karst.) selection forests at Siljansfors in Central Sweden. *Trees, Forests and People*, 12: 100392, 2023. ISSN 2666-7193. doi:https://doi.org/10.1016/j.tfp.2023.100392. URL https://www.sciencedirect.com/science/article/pii/S2666719323000249.

[5] Kajar Köster, Kaljo Voolma, Kalev Jõgiste, Marek Metslaid, and Diana Laarmann. Assessment of Tree Mortality After Windthrow using Photo-Derived Data. *Annales Botanici Fennici*, 46: 291–298, 08 2009. doi:10.5735/085.046.0405.

[6] Jarosław Paluch and Rafał Jastrzębski. Life histories of abies alba and picea abies growing in old-growth forests driven by natural gap-phase dynamics. *European Journal of Forest Research*, 142, 12 2022. doi:10.1007/s10342-022-01525-w.

[7] Giovanni Caudullo, Willy Tinner, and Daniele de Rigo. *Picea abies* in Europe: distribution, habitat, usage and threats. In J. San-Miguel-Ayanz, D. de Rigo, G. Caudullo, T. Houston Durrant, and A. Mauri, editors, *European Atlas of Forest Tree Species*, pages 114–116. Publications

Office of the European Union, Luxembourg, 2016. URL https://forest.jrc.ec.europa.eu/en/european-atlas/qr-trees/norway-spruce/. Joint Research Centre (JRC), European Commission.
